# Exon7 Targeted CRISPR-Prime Editing Approaches for *SMN2* Gene Editing in Spinal Muscular Atrophy (SMA) Disease Can Increase In Vitro SMN Expression

**DOI:** 10.1101/2022.03.21.484406

**Authors:** Sibel Pinar Odabas, Enes Bal, Gamze Yelgen, Ayse Simay Metin, Ebrar Karakaya, Gamze Gulden, Berranur Sert, Tarik Teymur, Yasin Ay, Nulifer Neslihan Tiryaki, Hasret Araz, Ilayda Cavrar, Cihan Tastan

**Affiliations:** Molecular Biology, Institute of Science and Technology, Üsküdar University, Istanbul, TURKEY; Transgenic Cell Technologies and Epigenetic Application and Research Center (TRGENMER), Üsküdar University, Istanbul, TURKEY; Molecular Biology and Genetics, Institute of Science and Technology, Kültür University, Istanbul, TURKEY; Molecular Biology and Genetics Department, Faculty of Engineering and Natural Science, Üsküdar University, Istanbul, TURKEY; HiDNA Biotechnology Inc., Istanbul, TURKEY

**Keywords:** CRISPR, Prime Editing, Spinal Muscular Atrophies, Gene Editing

## Abstract

Spinal Muscular Atrophy (SMA) is a fatal neuromuscular disease characterized by motor neuron loss and advanced muscle weakness, which occurs in functional SMN (Survival Motor Neuron) protein deficiency with *SMN1* gene-induced deletions and mutations. The incidence of SMA, which is an autosomal recessive disease, is 1/10,000 in the world. The SMN protein acts as a molecular chaperone in the formation of the spliceosome complex, which catalyzes the splicing of pre-mRNA, enabling mRNAs and non-coding RNAs to mature. Since the current *SMN1*-encoding Adeno-associated virus (AAV) or *SMN2* gene targeting antisense oligonucleotide-based strategies cannot provide long-term stable SMN expression in neuron cells, more effective methods need to be developed. CRISPR technology, which adds a new dimension to genetic engineering and gene therapies, makes it possible to treat many genetic diseases. In terms of SMA, some previous studies in the literature prove that it is possible to treat SMA with the CRISPR strategy. Homology Directed Repair (HDR)-based CRISPR technology, which results in a high rate of in-del (insertion-deletion) mutations rather than editing, was shown unsuitable for therapeutic applications. CRISPR-Prime editing (PE) technology is a new generation of gene editing approach that precisely provides various genomic modifications without the need for double-strand breakage or donor DNA sequences. CRISPR-Prime Editing method has also been used in rare diseases such as sickle cell anemia and Tay-Sachs, and their efficiency in editing various pathogenic mutations has been demonstrated. However, CRISPR Prime Editing-mediated gene editing for Spinal Muscular Atrophy (SMA) have not yet been investigated. The c.840 T-C transition and c.859 G-C transformations in the *SMN2* gene and the correction of these point mutations with a single pegRNA at the same time were investigated for the first time in this study. Here, we showed that CRISPR-PE systems could increase *SMN2* gene activity and SMN protein expression by ensuring exon 7 participation by editing c.840 T-C transition and c.859 G-C transformations. The fact that Prime Editing method showed the efficacy and stability of modifications in *SMN2* genes that were investigated in SMN-low Jurkat cells as a proof-of-concept. This study enabled the next step with the CRISPR-Prime Editing approach to be tested ex vivo in primary cell lines from SMA patients and SMN-low neuronal cells.

## Introduction

Spinal muscular atrophy (SMA) is a neuromuscular disease characterized by motor neuron loss and muscle atrophy resulting from a genetic defect. The incidence of SMA, which is an autosomal recessive disease, is 1/10,000 in the world. The SMN protein acts as a molecular chaperone in the formation of the splicosome complex, which catalyzes the splicing of pre-mRNA, enabling mRNAs and non-coding RNAs to mature (Bozorg Qomi et al., 2019). The factor that causes SMA disease is the loss of the ability to produce a full-length functional SMN protein as a result of various mutations and deletions in both alleles of the *Survival Motor Neuron 1* (*SMN1*) gene. The SMN protein, which is a housekeeping protein, plays a role in many vital processes for the cell such as snRNP biogenesis, transcription, translation, formation of stress granules, and transmission of cellular macromolecules (Singh et al., 2017). Failure to produce the SMN protein results in irreversible loss of anterior horn cells in lower motor neurons and brainstem nuclei in the spinal cord, resulting in further muscle weakness (Prior et al., 2009). There are two genes, *SMN1* and *SMN2*, which are associated with SMA disease and these genes express SMN protein (Chen et al., 2018). High diversity mutations in the *SMN1* gene that cause SMA disease (Wirth, 2000) result in exon 7 deficiency in the SMN protein in 95% of SMA patients. The critical region of 16 amino acids encoded by exon 7 is necessary for the oligomerization and stability of the SMN protein, and it enables the SMN protein to gain its functional property by folding (Lorson et al., 1999). In healthy individuals, the *SMN1* gene can produce 100% full-length (FL) SMN protein, while the *SMN2* gene can produce only ~10% FL SMN protein. The number of copies of *SMN2* in SMA patients correlates with the severity of the disease, and a low copy number of *SMN2* means the occurrence of more severe types of SMA (Bowerman et al., 2017). Many genetic diseases such as SMA in the world arise due to the formation of deletion, insertion, or base changes in the genome. At the beginning of the methods used in the treatment of SMA, the *SMN1* gene is given to the patient by using Adeno-associated viruses (AAV). (Zaidy and Mendel, 2019). AAV is a non-enveloped virus that can be modified to transfer DNA to target cells. Zolgensma (Zaidy and Mendel, 2019) treatment, which gives the *SMN1* gene to the patient using AAV serotype9, is not reliable in terms of providing long-term and high expression levels since it does not integrate into the cell genome (Lin et al., 2020). In recently published studies, data have been shown that the *SMN1* gene causes toxicity and lesion formation in tissues by transferring AAV vector systems to non-human primates (Verdera et al., 2020).

The technique that causes changes in the genome and allows the correction of these changes that underlie many diseases is known as the CRISPR/Cas9 gene-editing method (Jinek et al., 2012; Hsu et al., 2014; Yang et al., 2019). Some studies have observed that base modifications made in the SMN2 gene with the CRISPR system affect the increase in exon 7 participation in the *SMN2* gene. These studies show that SMA can be an ideal treatment model by proving the proof-of-concept with the CRISPR strategy (Zhou et al., 2018; Li et al., 2020; Lin et al., 2020). Prime Editing (PE), which has been recently defined as a new type of CRISPR, is a method that has a very low rate of secondary off-target mutation producing damage to the genome and can be used in all base changes (Cohen, 2019; Y. Liu et al., 2020). The CRISPR PE method results in higher efficiency and less indel formation compared to the Cas9-mediated Homology Directed Repair (HDR) method.

Point mutations can be repaired by HDR, but this method is not preferred in clinical studies due to the high percentage of off-target and low efficiency compared to the CRISPR PE method (Matsoukas, 2020; Schene et al., 2020). Prime editing guide RNAs (pegRNA) used in the CRISPR PE method allow many operations to be performed on the genome, including deletion, insertion, 12 base conversion, transversion, and transitions. The most important feature of the PE method, which is performed using pegRNA, is that it can perform gene editing in many combinations on the genome without creating a double-strand break in the DNA and without the need for a donor DNA template (Matsoukas, 2020). The strategy of Prime Editing, which has emerged as a versatile and sensitive genome editing method, is based on the combination of Cas9 nickase and reverse transcriptase (RT). The RT part of the Cas9-Reverse transcriptase protein uses the 18-20 nucleotide sequence on the pegRNA as a primer, takes the cut DNA as a template, and makes corrections on the newly synthesized DNA (Anzalone et al., 2019; Lin et al., 2020). Many gene-editing approaches in which CRISPR/Cas systems are used at the pre-clinical stage are available in the literature (Long et al., 2015; Tabebordbar et al., 2015; Yin et al., 2016; Bjursell et al., 2018; Maeder et al., 2019; Wu et al., 2019; Xu et al., 2019;). CRISPR Prime Editing (PE) method has been used in rare diseases such as sickle cell anemia and Tay-Sachs, and their efficiency in editing various pathogenic mutations has been demonstrated (Anzalone et al., 2019). In addition to increasing the reliability of gene therapy studies that can be done with the CRISPR-PE method, it is possible to cure point mutation-induced disease (ClinVar), which covers 50% of more than 75,000 known disease-related genetic variants in humans.

The most important change between *SMN1* and *SMN2* is the inhibition of exon 7 participation as a result of C-T transformation at 840 nucleotides in the *SMN2* gene. This nucleotide change causes the exonic splicing enhancer, which operates on exon 7, to be converted into a silencer. For this reason, the elements involved in the splicing phase cannot bind to the binding site and the production of SMN protein, which does not contain exon 7 at a rate of 80-95%, downregulates (Cartegni et al., 2006; M. A. Passini et al., 2011; Ross & Kwon, 2019; Schorling et al., 2020). Correct identification of exons by the splicing mechanism requires both classical splicing signals and multiple intronic and exonic cis-acting regulatory elements (Cartegni et al., 2002; Pagani et al., 2003). Exonic splicing regulatory elements are mainly recognized by trans-acting factors and are generally referred to as exonic splicing enhancers (ESEs) and exonic splicing silencers (ESSs). A nucleotide difference between *SMN2* and *SMN1* causes inefficient splicing of exon 7 in the SMN2 gene: the C-T transition at position +6 in exon 7 disrupts an SF2/ASF-ESE and/or creates an hnRNP A1-ESS (Cartegni et al., 2006; Kashima et al., 2007a; Kashima et al., 2007b; Vezain et al., 2010).

Due to mutations in the *SMN1* gene in SMA disease, functional SMN protein expression occurs only through the *SMN2* gene. Each individual, including healthy individuals, has the *SMN2* gene with a different copy number. Therefore, each SMA patient has at least one copy of *SMN2*. According to the data obtained, it has been determined that there is a relationship between the phenotypic severity of SMA patients and the *SMN2* gene copy number in the patients (Shorrock et al., 2018). It was observed that the phenotypic severity of the disease decreased with the increase in the *SMN2* copy number (Feldkötter et al., 2002; Shorrock et al., 2018).

In addition to the *SMN2* copy number, the higher the full-length SMN protein expression, the milder the disease phenotype (Feldkötter et al., 2002; Wirth et al., 2006). Likewise, it is known that the *SMN2* copy number has an important effect on the prolongation of the life span of patients, as well as its effect on the phenotype (C. C. Wang et al., 2009). Since the *SMN2* gene is common in patients with all SMA types, it is aimed to increase SMN protein expression by making modifications with CRISPR Prime Editing, an innovative gene-editing method over the *SMN2* gene. In addition, it is planned to investigate a new generation gene modification strategy that can be adapted to all SMA types in this way. In studies conducted to explain the rare phenotype that changes independent of *SMN2* copy number, a positive modifying effect of the c.859 G-C natural variants for SMA disease has been shown (Prior et al., 2009; Vezain et al., 2010; Reed and Zanoteli, 2018). The *SMN2* c.859 G-C (p.Gly287Arg) variant has been proven in case studies to increase exon 7 involvement, thereby reducing the phenotypic severity of the disease due to an increase in the amount of full-length SMN protein (~20%) (Prior et al., 2009; Yépez et al., 2020). Studies have shown that this variant does not affect the binding of HTra2β or create a new SF2/ASF enhancer. This variant has been shown to increase exon 7 involvement due to the degradation of an hnRNP A1-dependent silencer and binding domain (Vezain et al., 2010). The incidence of this variant is 1/150 in healthy individuals, while it is 2/375 in individuals with SMA (Bernal et al., 2010; Vezain et al., 2010).

Investigations of CRISPR PE-mediated gene editing in Spinal Muscular Atrophy (SMA), a rare and genetic disease, have not yet been conducted. In this study, we have targeted modifications of *the SMN2* gene by using CRISPR PE technology, which has many advantages in terms of gene editing, since it is possible to increase functional SMN protein expression by correcting only 2 points (840. T-C and 859. G-C) mutations. With this study, for the first time, the CRISPR-PE strategy, which has proven superiority in terms of target finding, modifying specific nucleotides, and reliability, is used in the treatment of SMA disease. By examining the feasibility of this strategy, a proof-of-concept was provided for the first time specifically for SMA.

The main goal of this project, which includes gene therapy strategies, is to ensure that the full-length functional SMN protein expression level is kept high and stable in the long term by manipulating the splicing process. The modifications determine the inclusion-skipping of the exon 7 region in *SMN2* gene that plays a key role in the function of the SMN protein and making it permanent with the CRISPR-PE system. This study was a proof-of-concept study in Jurkat cells that produce low levels of SMN protein, edited with CRISPR-PE and 3 different pegRNA aiming to modify two different nucleotides in the *SMN2* gene. This study let us screen fifteen other mutations on the gene to determine the most efficient SMN protein up-regulation in primary SMA type1 fibroblast cells and neuron cell line A172 having a low level of SMN protein.

## Materials & Methods

### Designing pegRNAs for the CRISPR Prime Editing method

The database https://primeedit.nygenome.org/ was used for the design of prime editing guide RNA (pegRNA) on the *SMN2* gene. Table 1 shows the design of pegRNA1 for c.859 G-C transformations for target sites, pegRNA2 for c.840 T-C transition, and pegRNA3 including both target sites. For the modification of target sites, reverse transcriptase (Reverse transcription template, RTT) and primer binding sequences (Primer Binding Site, PBS) were designed (Table 1). The pegRNA synthesis sequences from the pU6 promoter to the elongation sequence are shown (Table 2). The designed pegRNA sequences were synthesized with the pU6 promoter by GenScript and cloned into the plasmid pHIV-EGFP (Addgene, #21373).

**Table 1.**
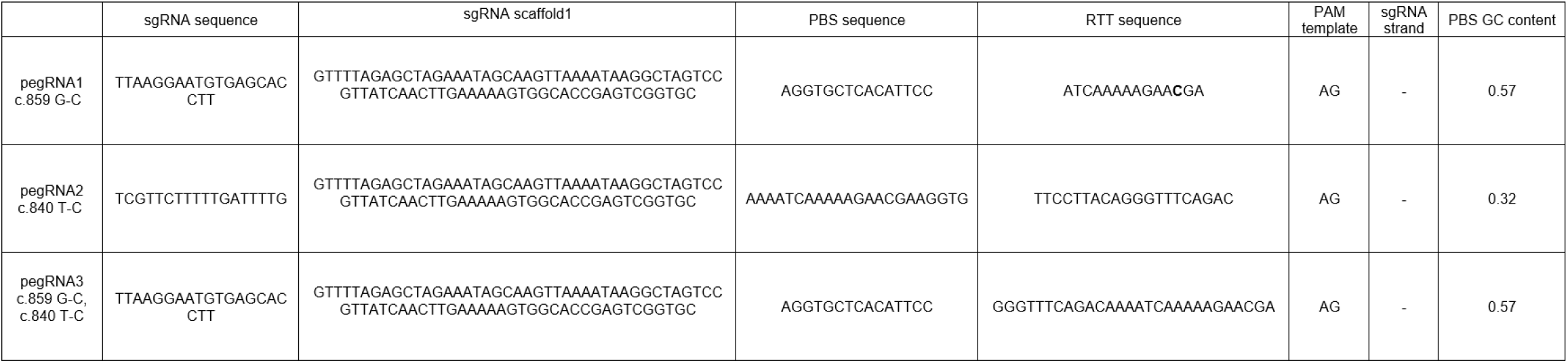
pegRNA designs

**Table 2.**
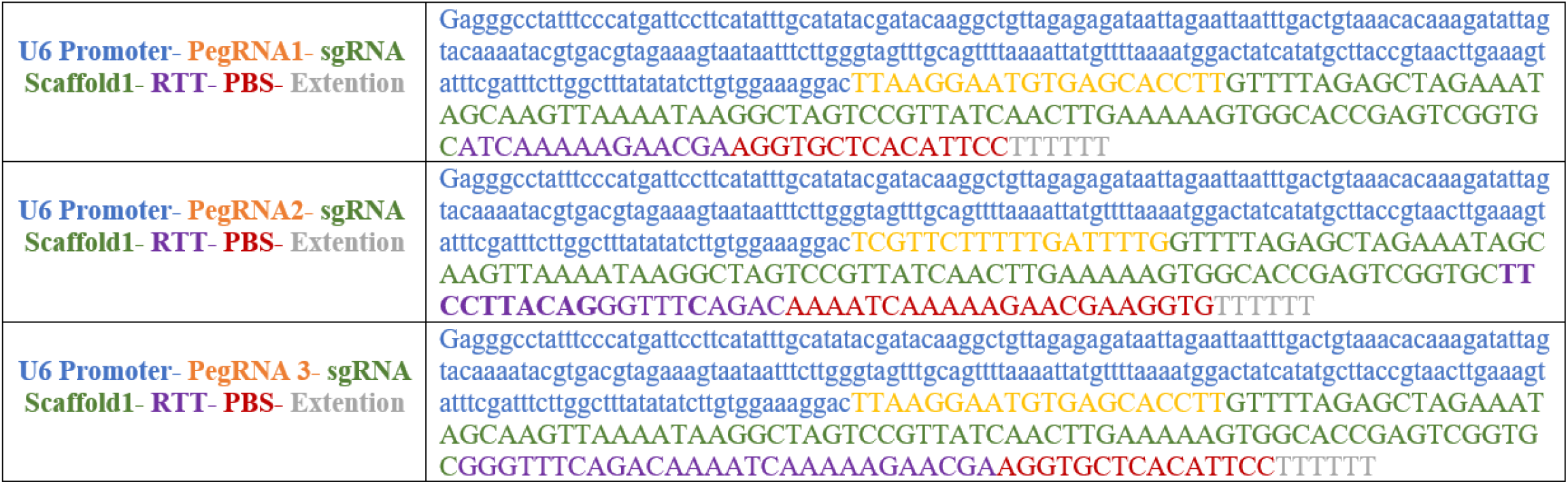
pegRNA synthesis sequences

### Transformation and Isolation

Bacteria of Competent *E. coli* DH5α strain (NEB® 5-alpha Competent *E. coli* (High Efficiency) NEB C2987H) were cultured in a liquid LB medium at 37°C. The bacterial strain was stored at −80°C as stock in LB medium with added glycerol. Bacteria were transformed with plasmid DNAs using the heat shock method according to the manufacturer’s instructions. Bacteria were inoculated on LB agar plates with ampicillin and incubated at 37°C overnight. Plasmid isolation was performed from selected bacterial colonies with the CompactPrep Plasmid Maxi Kit (#12863, QIAGEN). The following quality control steps for DNA and virus production have been evaluated by considering the specified acceptance criteria. DNA concentration was measured in ng/μl with Microplate ELISA Reader (FLUOstar Omega) and it was evaluated whether the purity value was between 1.8<A260/A280<2.0. For the control of the isolated plasmids, DNA samples loaded on the gel prepared in 1X TAE buffer solution using 1.5% agarose gel electrophoresis system were run at 90 V and 60 minutes, and plasmid DNA samples with <1% of bacterial DNA contamination were used for lentivirus production.

### Lentivirus Production

The designed 3 pegRNA sequences were synthesized with the pU6 promoter by GenScript and cloned into the lentiviral pHIV-EGFP (Addgene, #21373) plasmid. For lentivirus production, lentiviral pLenti-PE2-BSD plasmid DNAs encoding CRISPR-PE containing the blasticidin resistance gene, control pHIV-EGFP non-coding pU6-pegRNA1/2/3, and coding GFP and pHIV EGFP coding pU6-pegRNA1/2/3 and GFP were transfected with pSPAX2 and pVSV-G envelope and packaging plasmid DNAs. After all plasmids (2:1 pSPAX2: 1 pVSV-G) were treated with P-PEI (Polyethylenimine) transfection reagent (Merten et al., 2016), lentivirus production was performed using the host cells, HEK293T, as in our previous studies (Chen et al., 2018; Tastan et al., 2018; Tastan et al., 2020). Packed recombinant lentiviruses were obtained by collecting HEK293T cell supernatants 72 hours after transfection. Produced lentiviruses were concentrated 100x with Lenti-X Concentrator (#631232, Takara) to increase virus concentration.

### Lentivirus Titration

For the titration of lentiviruses, titration steps in a previously published study were performed (Taştan et al., 2020). Jurkat (RPMI medium containing 10% fetal bovine serum, 200 U/ml penicillin/streptomycin antibiotic, 2 mM L-glutamine, 1X MEM vitamin solution, and 1X NEAA) cells were seeded in 96-well petri dishes in 100 μL medium. Wells were adjusted to contain 100x-concentration lentivirus at 10μL, 3μL, 1μL, 0.3μL, 0.1μL, and 0.03μL. 50 μL of lentivirus dilution from each concentration was transferred to the wells containing Jurkat cells, respectively, and the total volume was adjusted to be 150 μL. Afterward, the cells were incubated for 3-4 days at 37°C, %5 CO_2_. After the cells were transferred to 96-well plates, Jurkat cells containing pLenti-PE2-BSD (Addgene, #161514) were treated with 10 μg/ml blasticidin for selection of blasticidin and incubated for 72 hours. Afterward, cell viability was tested by staining with a 7-AAD antibody in flow cytometry. Finally, cells encoding CRISPR-PE with >99% blasticidin resistance were selected and cultured. Titration of pegRNA-GFP and control GFP lentiviruses was calculated by measuring the GFP expression level in flow cytometry. Subsequently, 3 pegRNA-GFP or control GFP lentivirus were individually transduced into CRISPR-PE (+) Jurkat cells at a rate of 5 MOI.

### Cell Culture

SMA Type 1 fibroblast cells (Coriell Institute, GM00232, EMEM medium with 15% fetal bovine serum (FBS), 200 U/ml penicillin/streptomycin antibiotic, 2 mM L-glutamine and 1X NEAA), HEK293T (Gibco DMEM medium with 10% FBS and 1% penicillin/streptomycin, L-Glutamine medium), HeLa (Gibco DMEM medium with 10% FBS and 1% penicillin/streptomycin, L-Glutamine), A172 (DMEM medium with 15% FBS, 200 U/ml penicillin/streptomycin antibiotic, 2 mM L-glutamine), SH-SY5Y (DMEM medium with 15% FBS, 200 U/ml penicillin/streptomycin antibiotic, 2 mM L-glutamine, 1% sodium pyruvate), U87 (DMEM medium with 15% fetal bovine serum, 200 U/ml penicillin/streptomycin antibiotic, containing 2 mM L-glutamine) and HMC3 (MEM medium with 15% FBS, 200 U/ml penicillin/streptomycin antibiotic, 2 mM L-glutamine, 1% sodium pyruvate) were cultured. Cells were incubated at 37°C, 5% CO_2_.

### Transfection of AAV vector containing the *SMN1* gene

The *SMN1* AAV Vector (Human; hSyn; Luc) (#444551011210) containing the functional, synthetic *SMN1* gene (GenBank accession code, NM_000344) was obtained from Applied Biological Materials Inc. *SMN1* AAV plasmid DNA was transferred to SMA fibroblast cells for SMN expression control after treatment with PEI (Polyethylenimine) transfection reagent (Merten et al., 2016). SMN expression was then analyzed by flow cytometry.

### Flow Cytometry

SMA fibroblast, Jurkat, HEK293T, HeLa, and neuronal cell lines A172, SH-SY5Y, U87, and HMC3 cells were incubated with anti-SMN-AlexaFlour 647 (Santa Cruz Biotechnology, Anti-SMN Antibody (2B1)) monoclonal antibody to determine SMN expression levels. Fixation/permeabilization procedures were performed using the BDCytofix/Cytoperm Fixation/Permeabilization Kit (Cat. No. 554714). After the cells were collected to form 100,000 cells, 200 μl of Fixation/Permeabilization solution (BD) was added and incubated at 4C for 20 minutes. After washing with BD Perm/Wash™ Buffer (BD). cells were stained with anti-SMN-AlexaFlour 647 at a dilution ratio of 1:50 and incubated at 4C for 30 minutes. Cells were washed with BD Perm/Wash™ Buffer (BD) and analyzed by flow cytometry (Beckman Coulter, CytoFlex) to assess the percentage and the Mean Fluorescence Intensity (MFI) level of SMN expression.

### Cell Survival Analysis

Survival analysis was performed to control the cell viability of Jurkat cells. Trypan blue (Biological Industries, #03-102-1B) was applied to identify and count surviving cells. Cell counting and viability analysis were performed with the BIO-RAD TC20 Automated Cell Counter.

### Statistical Approach

Data recorded by flow cytometry were analyzed using CytExpert (Beckman Coulter). Statistical analyses were performed using GraphPad Prism 8.0.1 software (GraphPad Software, Inc., San Diego, CA, USA) with a two-tailed t-test using independent mean values. Error bars represent Standard Error of Means (SEM). For all experiments, significance was defined as p<0.05 and *NS*= Non-Significant.

## Results

To determine the potential targets in the *SMN2* gene for the study, we first focused on the variant analyses and site-specific mutagenesis studies reported in the literature (Cartegni et al., 2006; Kashima et al., 2007a; Kashima et al., 2007b; Prior et al., 2009; Vezain et al., 2010; Reed and Zanoteli, 2018). Here, we aimed first to perform position-specific editing of the nucleotides including (840. T-C and 859. G-C) in exon 7 associated with the exon 7 skipping mechanism as a result of *SMN2* pre-mRNA processing (Cartegni et al., 2006; Prior et al., 2009), using the novel gene-editing approach CRISPR-PE (**Figure 1a**). To test the efficacy of the pegRNAs to edit the points, we set up an experimental proof-of-concept strategy in the SMN-low Jurkat cell line (**Figure 1b**). This will let us determine the specific positions capable of increasing the SMN protein through *SMN2* gene expression.

**Figure 1:**
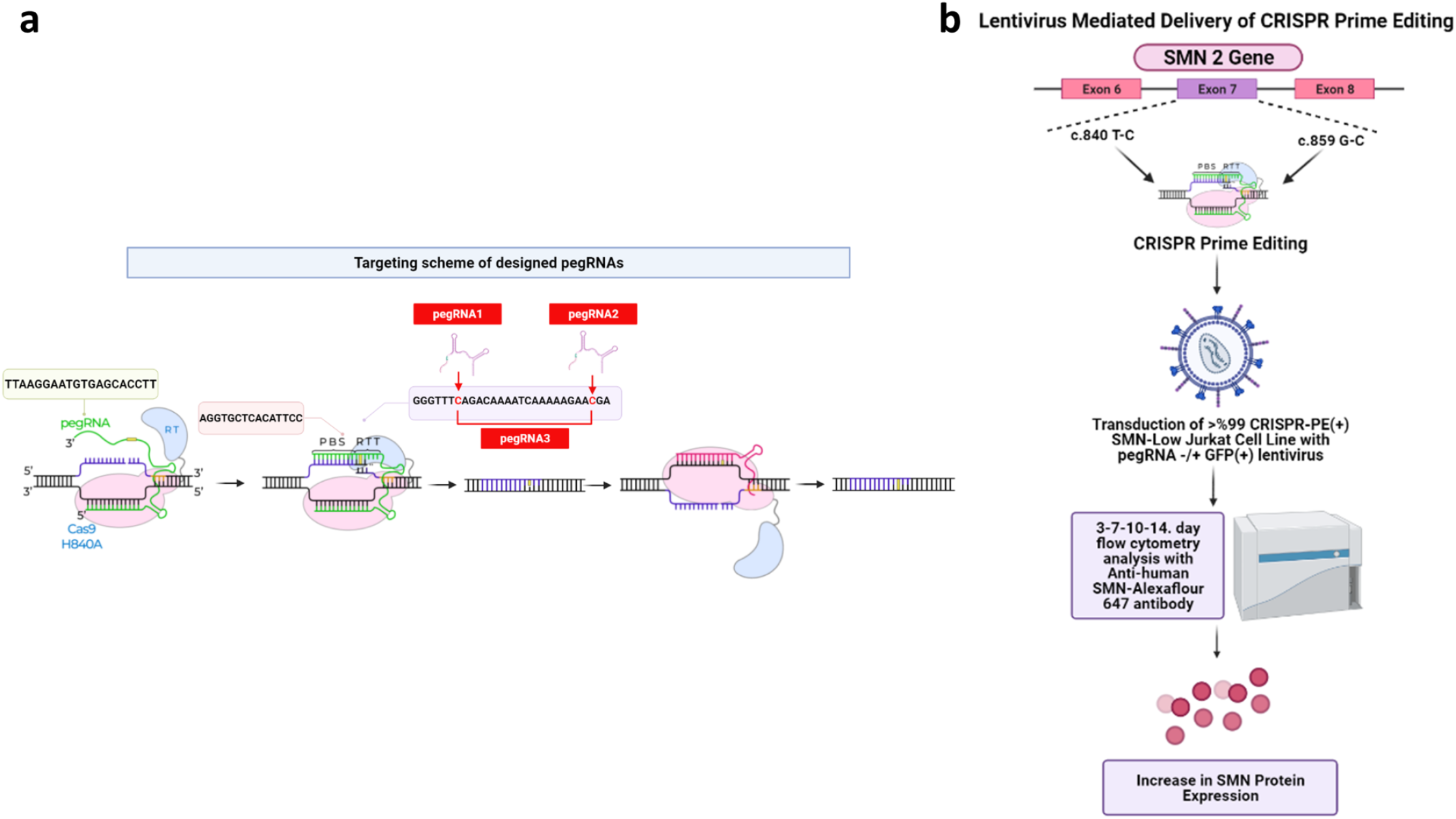
*SMN2* Gene Editing and Experimental Proof-of-Concept Strategy in SMN-low Jurkat cell line. **a**. Modifications of *the SMN2* gene by using CRISPR-PE technology by correcting the points (840. T-C and 859. G-C) mutations. pegRNA1 to modify 859. G-C, pegRNA2 to modify 840. T-C, and pegRNA3 to modify both nucleotides. **b**. Experimental strategy to assess upregulation of SMN protein expression in Jurkat cell line. The cells were first transduced with CRISPR-PE and Blasticidin resistant gene coding lentivirus to develop >%99 CRISPR-PE (+) Jurkat cell line. Next, PE(+) Jurkat cells were transduced with pegRNA(+/−) GFP coding lentivirus. The cells were analyzed using flow cytometry after fixed and stained with the human anti-SMN monoclonal antibody at the 3, 7, 10, and 14th days of second transduction.

To develop CRISPR-PE (+) Jurkat cells, the cell line was firstly transduced with pLenti-PE2-BSD lentivirus and selected with Blasticidin. Next, the pegRNAs under the pU6 promoter in pHIV-EGFP lentiviruses were used to transduce the PE(+) Jurkat cells (**Figure 2a**). pegRNA expressing cells after transduction in the Jurkat cells (**Figure 2a**) and after transfection into the HEK293 cell line (**Figure 2b**) was determined by GFP expression using flow cytometry and fluorescence microscopy, respectively. Afterward, SMN upregulation in the pegRNA(+) PE(+) Jurkat cells were determined by anti-SMN monoclonal antibody at day 3, day 7, and day 10 post-infection (**Figure 2c**). Independent analyses performed on day 7 and day 10 showed that SMN expression increased from <10% in control GFP positive pegRNA-negative Jurkat cells to >15% in pegRNA1 and ~15% in pegRNA3 expressing Jurkat cells **(Figure 2c)**. We determined control GFP(+) PE(+) Jurkat cells did not show an increase of SMN level; however, pegRNA1 and pegRNA3 GFP (+) PE(+) Jurkat cells could upregulate SMN expression after 7 days of the transduction (**Figure 2c**). Interestingly, it was observed that SMN expression completely decreased to 0% at day 10 in the PE(+) Jurkat cells encoding pegRNA2 providing 840. C-T exchange (**Figure 2c**). pegRNA2 GFP(+) PE(+) Jurkat cells showed an increase in MFI level of SMN at day 3 but the cells did not survive after day 10 (**Figure 2c**). These results showed that SMN expression by the modification of the *SMN2* gene is affected by the edited nucleotide.

**Figure 2:**
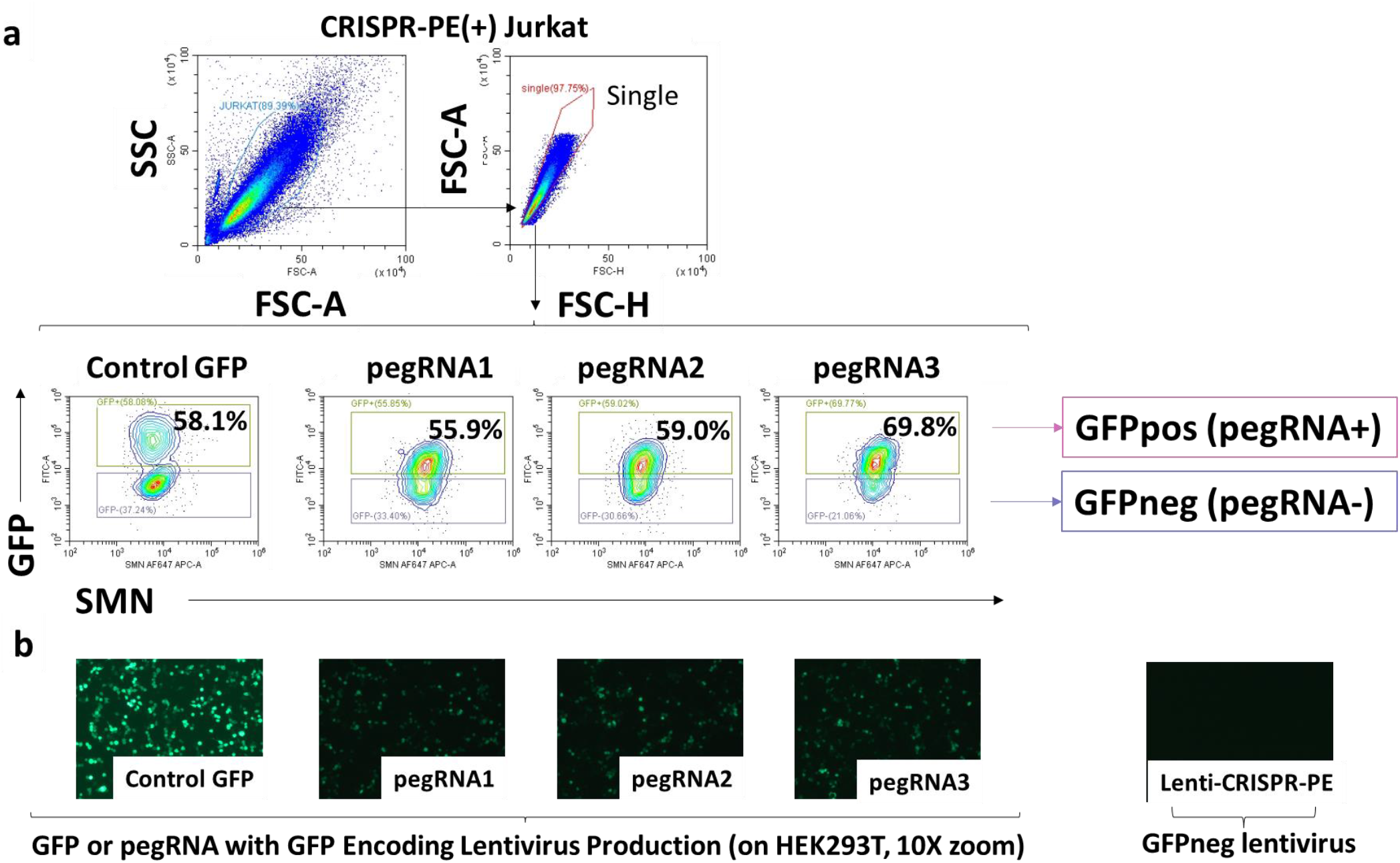
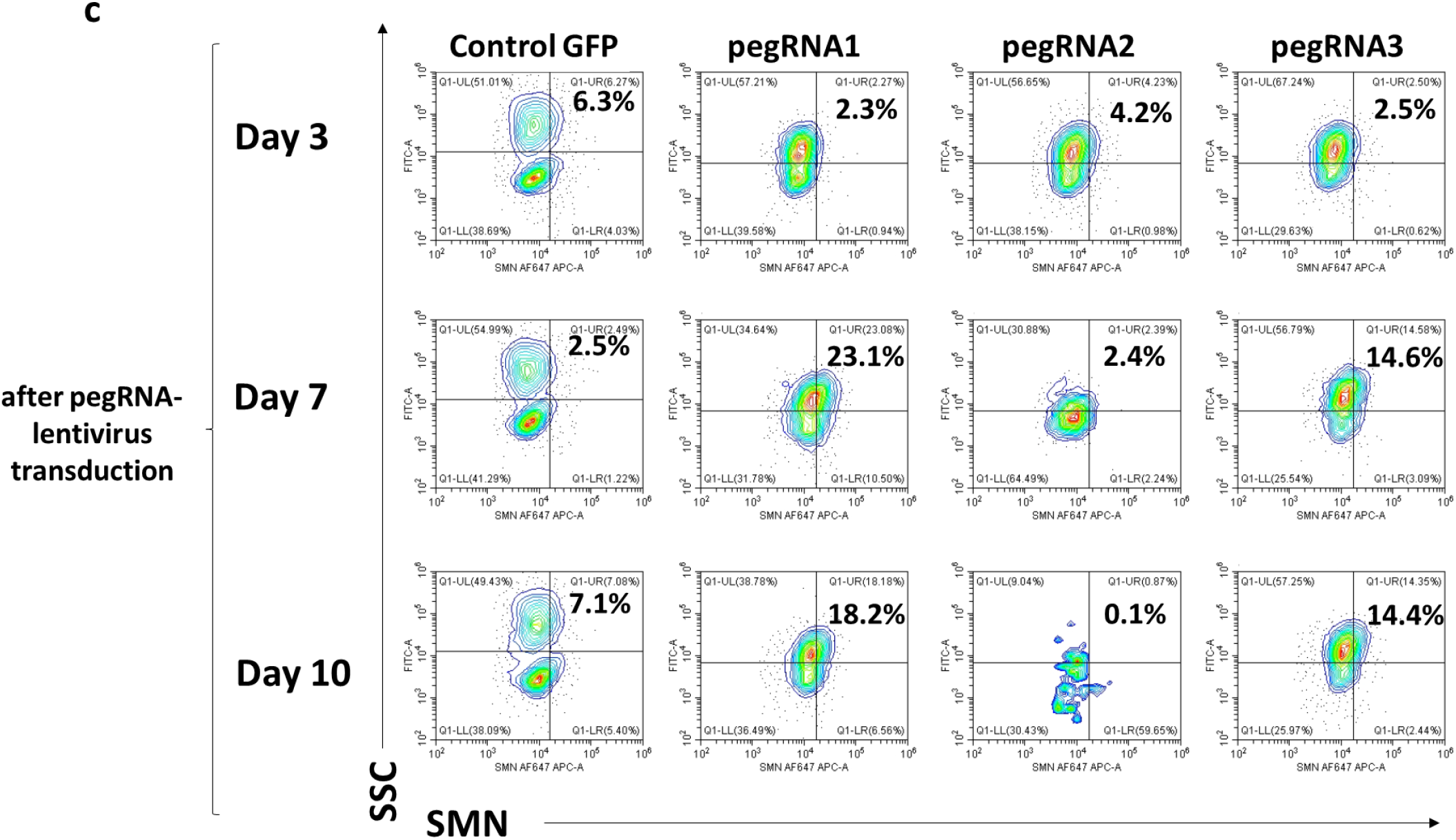
The effect of gene modification with CRISPR-PE and pegRNA system on SMN expression in SMN-low Jurkat cells. **a**. Fixed Jurkat cells were stained with anti-human SMN Alexa Flour 647 monoclonal antibody and analyzed by flow cytometry. PE(+) Jurkat cells expressing pegRNAs were determined to be GFP positive. PE(+) Jurkat cells transduced with GFP-encoding lentivirus as the pegRNA negative control. **b**. Production of the control GFP positive lentiviral vector and pegRNA-GFP encoding lentiviral vectors on HEK293T cells was demonstrated by a fluorescent microscope at 5X. **c**. Upregulation of SMN protein expression in PE positive Jurkat cells after transduced with control GFP or pegRNA-GFP encoding lentiviruses at days 3, 7, and 10.

Next, we wanted to assess the increase of SMN expression by quantifying mean fluorescence intensity (MFI) concerning the time after transduction of the pegRNAs. First, the SMN MFI values of GFP-expressing and non-GFP-expressing cells were analyzed separately (**Figure 3a**). The background SMN-low levels in Jurkat cells were determined based on the SMN MFI level in the PE non-coding pegRNA transduced Jurkat cells. Next, the SMN MFI values in GFP-expressing and non-GFP-expressing cells were subtracted from each other, and the increase in SMN MFI in pegRNA-transduced PE(+) Jurkat cells was considered as the true CRISPR-PE effective increase (**Figure 3b**). SMN MFI level in pegRNA (+) GFP (+) PE (+) Jurkat cell were determined statistically significant comparing to pegRNA (+) GFP (+) PE (−) and pegRNA (−) GFP (+) PE (+) Jurkat cells (**Figure 3b**). The level of SMN MFI in pegRNA (+) GFP (+) PE (−) Jurkat cells was determined as the baseline SMN MFI difference, and this baseline difference level was then subtracted from the SMN MFI difference value in pegRNA (+) GFP (+) PE (+) Jurkat cells (**Figure 3c**). The increased SMN expression in pegRNA and PE encoding cells was determined as the CRISPR-PE system effect. According to this analysis, statistically significant SMN upregulation was detected in pegRNA1 and pegRNA3 transduced PE (+) Jurkat cells after day 7 (**Figure 3c**). These results showed that the pegRNA 1 and pegRNA3 designs could increase SMN expression with the CRISPR-PE system.

**Figure 3:**
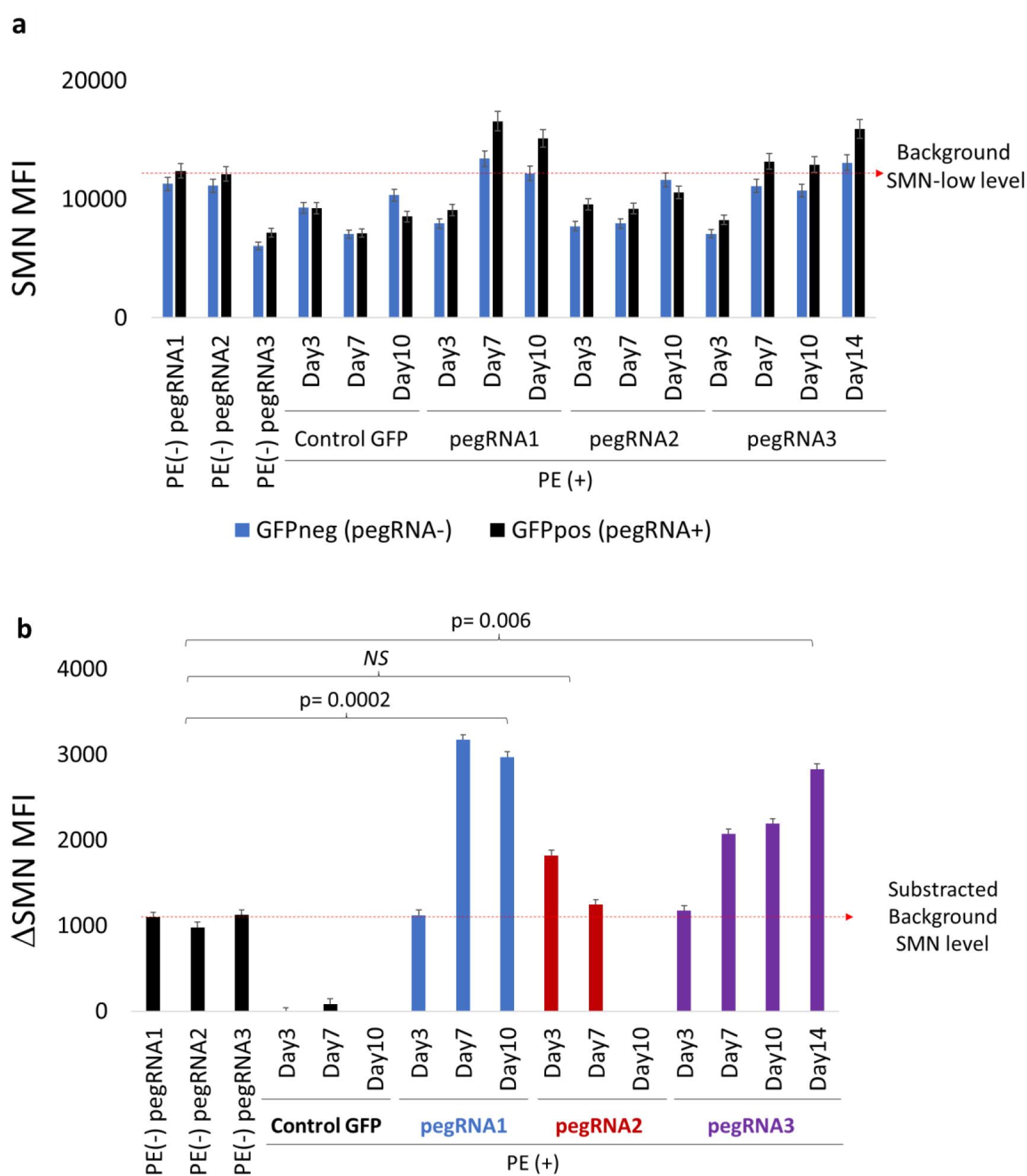
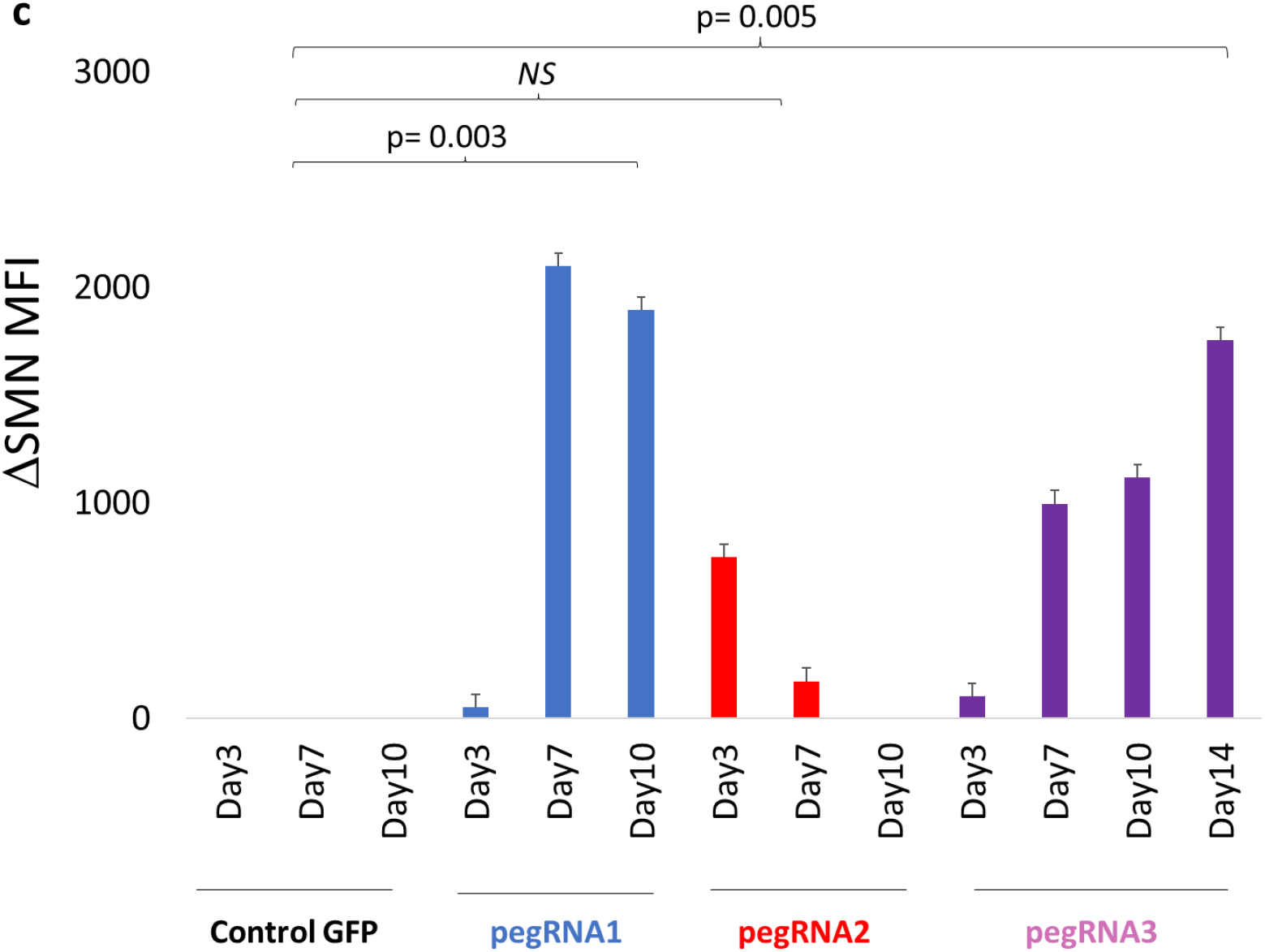
Upregulation of SMN protein expression determined as MFI level. **a**. Bar graph showing SMN MFI values of GFP-expressing and non-GFP-expressing cells in the Jurkat cells that are controls including pegRNA (+) GFP (+) PE (−) and pegRNA (−) GFP (+) PE (+), and pegRNA (+) GFP (+) PE (+). The red arrow shows background SMN-low level in Jurkat cells based on the SMN MFI level of the PE non-coding pegRNA transduced Jurkat cells. **b**. The bar graph shows SMN MFI values of the cells after subtraction of the SMN MFI values of GFP-expressing and non-GFP-expressing cells. The red arrow shows subtracted SMN MFI level in pegRNA (+/−) GFP (+) Jurkat cells based on the SMN MFI level of the PE (−) pegRNA (+) Jurkat cells. **c**. The bar graph shows the increased SMN expression in pegRNA (+) and PE (+) cells after subtraction of the level of SMN MFI in pegRNA (+) GFP (+) PE (−) cells from the SMN MFI level in pegRNA (+) GFP (+) PE (+) Jurkat cells. p< 0.05. NS= Non-Significant.

To analyze the persistence of SMN protein expression with the CRISPR-PE system, PE (+) Jurkat cells expressing pegRNA3 provide modification in both target regions (840. T-C and 859. G-C) were cultured for 14 days and SMN expression was determined (**Figure 4a**). It was shown that stable SMN protein production was maintained for 14 days and had a statistically significant increase compared to pegRNA3 (+) non-PE-expressing and GFP (+) Jurkat cells expressing PE but not pegRNA (**Figure 4b and 4c**). These results showed that the CRISPR-PE approach can provide single nucleotide changes on the *SMN2* gene that can permanently increase SMN expression.

**Figure 4:**
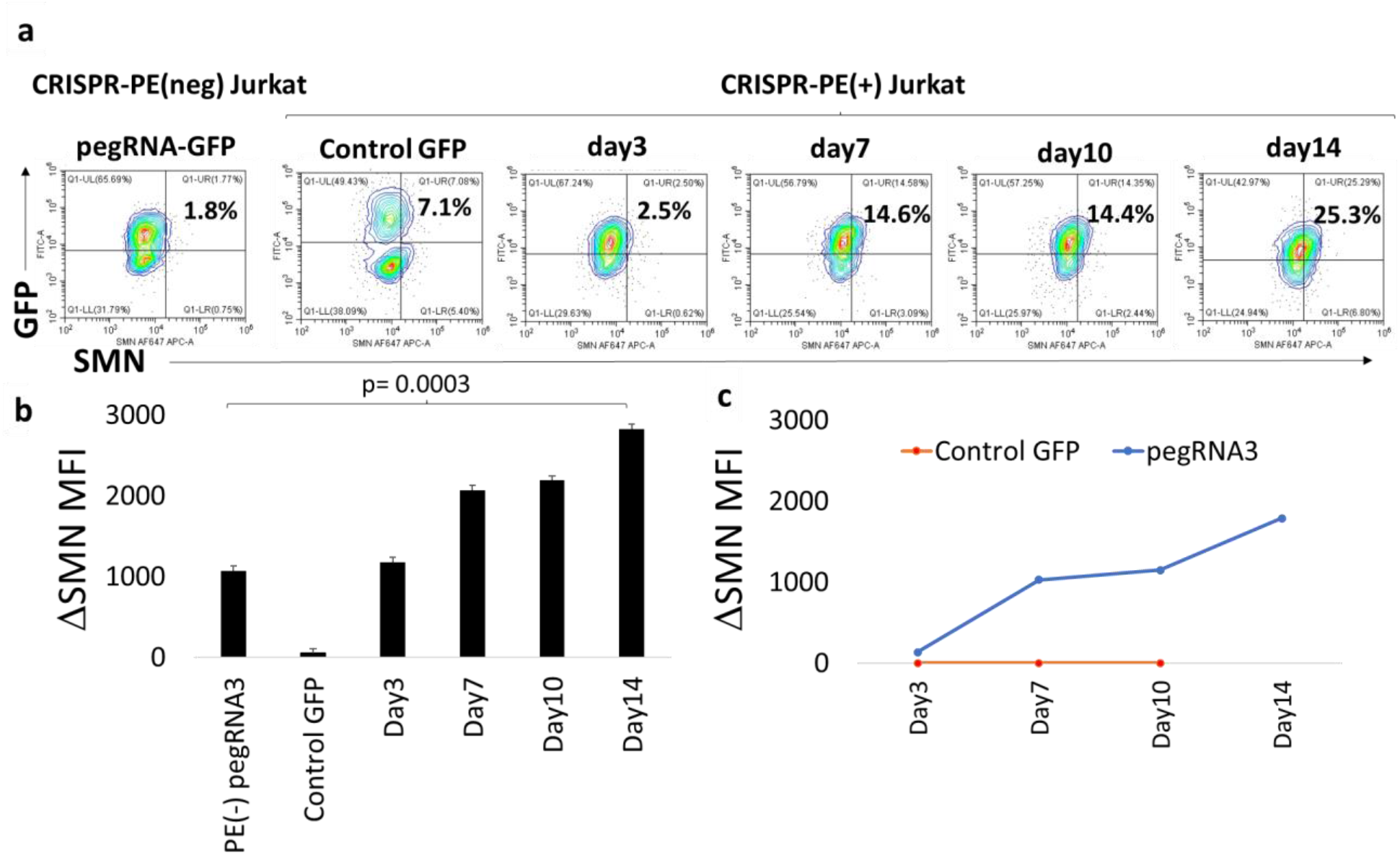
Time-dependent manner of SMN upregulation in pegRNA3 (+) PE(+) Jurkat. **a**. Flow cytometry plots show GFP and SMN expression of the cells that were either pegRNA (+) GFP (+) PE (−), pegRNA (−) GFP (+) PE (+), and pegRNA (+) GFP (+) PE (+) analyzed at day 3, 7, 10, and 14. **b**. Bar graph shows SMN MFI level of GFP (+) cells after subtracted from the SMN MFI level of GFP (−) cells in the same culture. **c**. The graph shows SMN MFI upregulation in pegRNA3 (+) GFP (+) PE (+) Jurkats in a time-dependent manner with respect to the pegRNA (−) GFP (+) PE (+) cells after subtracted from the SMN MFI difference level of pegRNA3 (+) GFP (+) PE (−) cells. p< 0.05. NS= Non-Significant.

Afterward, we wanted to determine SMN-low primer cells or neuronal cell lines to test with the CRISPR-PE system along with pegRNA1 and pegRNA3 which we determined to be effective on Jurkat cells. In our study, the primary fibroblast cell line taken from an SMA type 1 patient, was first obtained from the Coriell Institute. Then, in parallel, SMN expression in HEK293T and HeLa cell lines was assessed with anti-SMN monoclonal antibody (**Figure 5a and 5b**). The plasmid DNA including the human synapsin1 promoter controlling *SMN1* gene expression from Applied Biological Materials Inc (*SMN1* AAV Vector (Human; hSyn; Luc)) was transfected into the primary SMA fibroblast cell line to determine the capacity of the cells for SMN upregulation. Thus, we have shown that the expression level of SMN in fibroblast cells can be increased to a level similar to the level of SMN expression in HEK293T and HeLa cells (**Figure 5a**). SMN expression (~10%) in the primary SMA fibroblast cells can be upregulated with transgene *SMN1* upregulation, showing upregulation of the SMN expression over transfection of the transgenic *SMN1* gene up to a level (~70%) (**Figure 5a and 5c**). This suggests that the primer fibroblast cells isolated from SMA type1 diagnosed patients have capable of physiological SMN expression. Next, we wanted to determine SMN protein levels in neuronal cell lines including A172, SH-SY5Y, U87, and HMC3. Among these cell lines, the A172 glioblastoma cell line had a low SMN expression level and can be used in further experiments with CRISPR-PE (**Figure 5d**). These findings showed that A172 and primer SMA fibroblast cells can be used as a cell line with low SMN expression suitable for performing *SMN2* gene-targeted CRISPR-PE activity screens.

**Figure 5:**
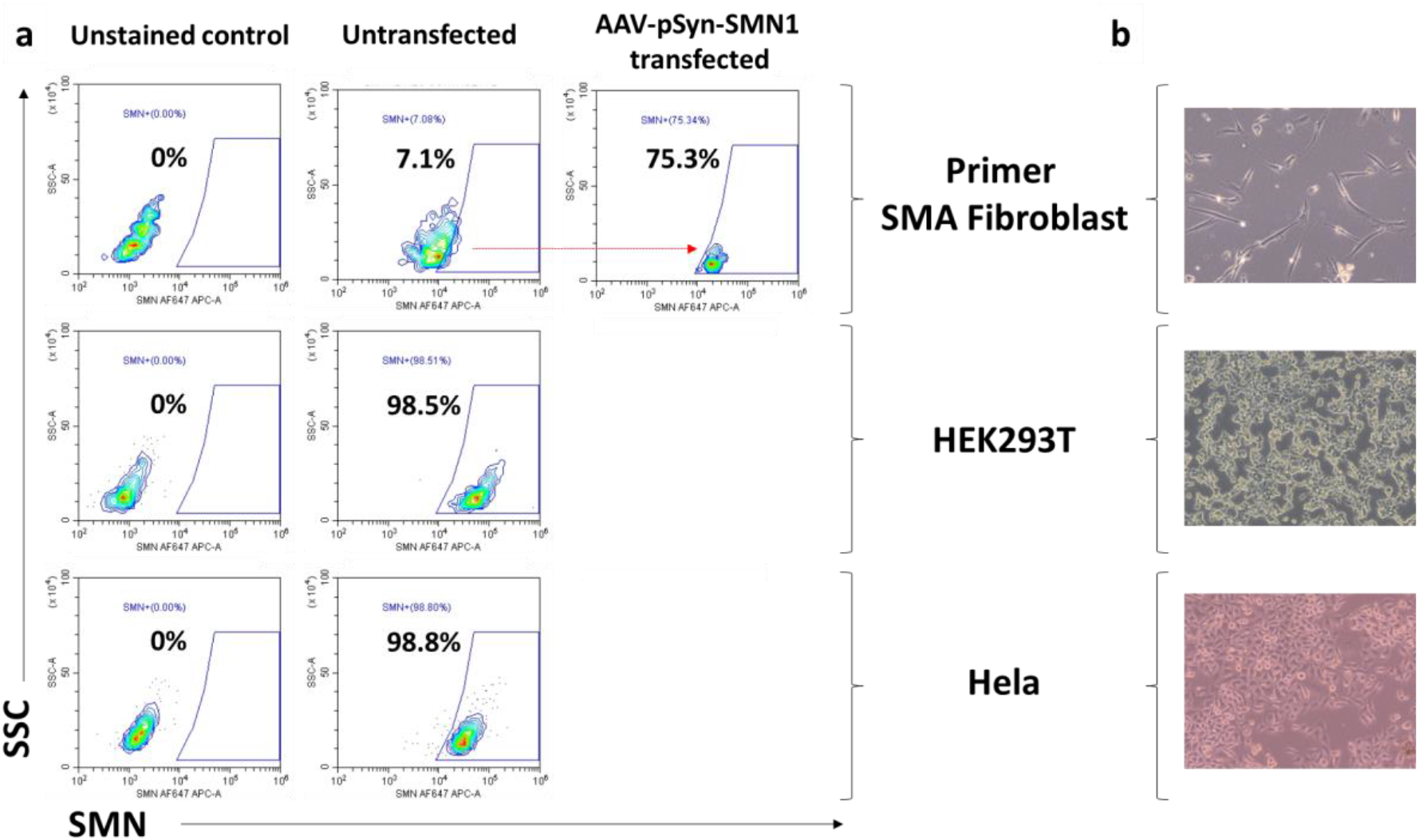
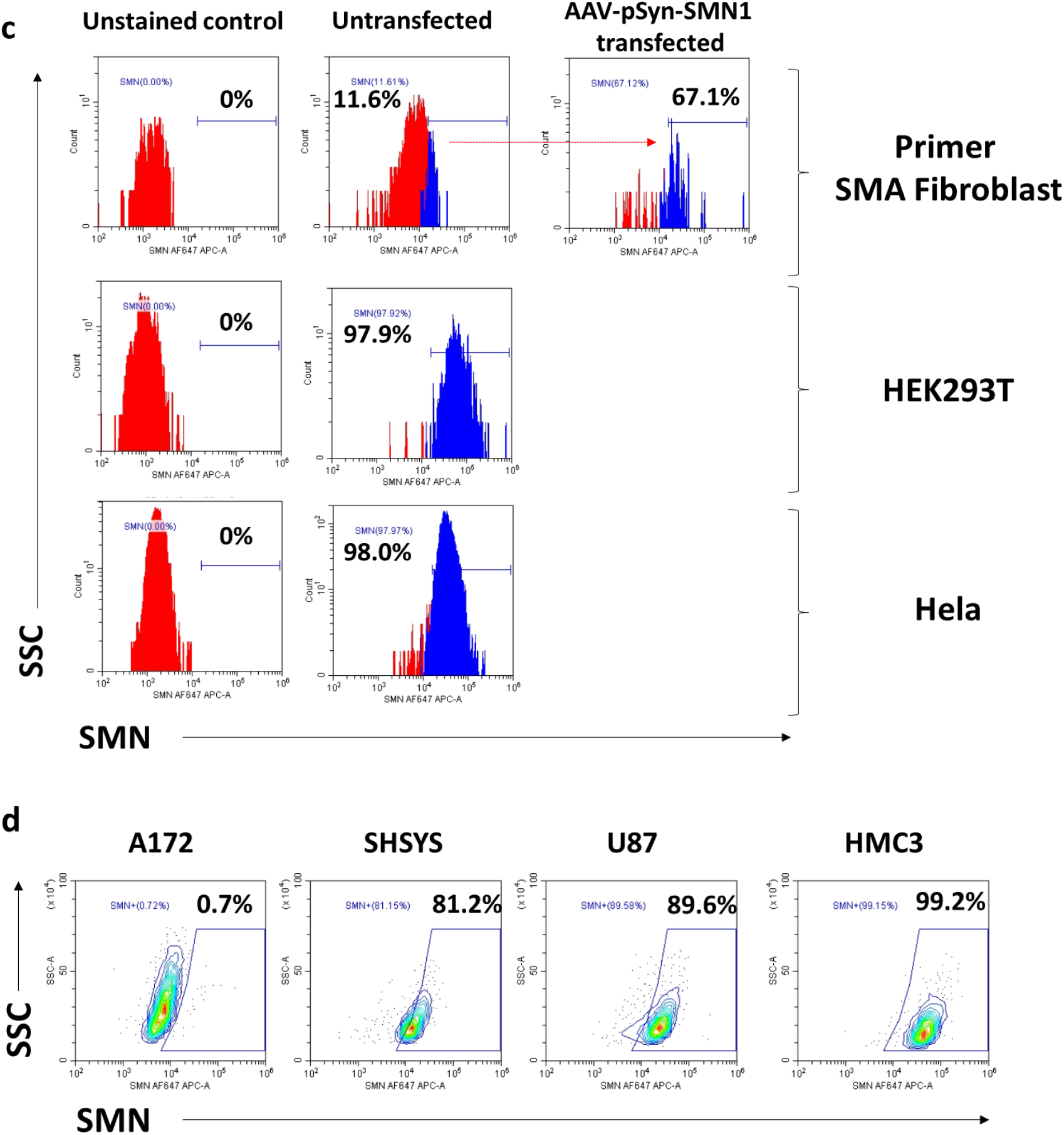
Determination of SMN-low primer fibroblast and neuronal cell lines for further CRISPR-PE studies. **a**. As a result of transgenic *SMN1*-containing AAV plasmid DNA (Human; hSyn; Luc) transfer process induced by synapsin 1 promoter in isolated fibroblast cells from primary SMA type1 patient, SMN expression was measured after 24 hours using Anti-Human SMN-AlexaFlour 647 monoclonal antibody by flow cytometry method. **b**. Imaging of SMA fibroblast, HEK293, and Hela cell lines under the light microscope at 5X. **c**. Histograms of SMN expression in SMA model primer fibroblast, HEK293, and Hela cell lines. **d**. SMN expression levels of neuronal cell lines.

The *SMN2* gene was planned considering the mutagenesis studies in exon 7/intron 7 regions, and it was seen that the base changes created in these two regions greatly affected the increase in exon 7 participation (**Figure 6**). Optimum pegRNA sequences that perform target-directed base modification with high sensitivity and provide high efficiency in *SMN2* gene exon 7 participation increase will be examined and determined within the scope of this work package. Within the scope of this project, it is aimed to use the CRISPR-PE system prepared to achieve a high on-target score and low off-target score in motor neuron cells forthe *SMN2* gene, and to determine pegRNA designs that can keep functional SMN protein expression high and stable in the long term. In the continuation of this pioneering work, we aim to make permanent nucleotide changes on the *SMN2* gene with CRISPR-PE technology and to prevent exon 7 skipping that occurs during pre-mRNA processing by determining the optimum pegRNA designs that make this change specific to motor neuron cells. With the participation of exon 7 in the protein production process, it is aimed to increase the expression of functional SMN protein and to keep this expression increase stable in the long term. In the continuation of our study, *SMN2* region-specific mutagenesis studies and chemical structure analyses made to increase *SMN2* gene participation were examined and specific positions and nucleotide intervals that could increase exon 7 participation were determined. Accordingly, the *SMN2* gene will provide base changes in exon 7, A11G, A7C, A51C, T44G, A54G, 40C54G, *SMN2*.6G11G, SMN2.6A11G, *SMN2*.10G11G, and intron 7, G100A, and 10 in intron 7. PegRNA sequences were designed to delete bases at positions 290 and 290 (C and G) and GCAGAC bases between positions 290-295 (**Figure 6**). We will continue to scan with the CRISPR-PE system that will ensure these changes. At the end of this stage, pegRNA sequences with high on-target and low off-target effects will be determined at targeted positions to increase *SMN2* gene participation. We will examine the variation of SMN expression level in heterogeneous cell populations as a result of a transfer of lentiviruses encoding CRISPR-PE in Jurkat, SMA type 1 stable-fibroblast, and SMN-low A172 neuronal cells to observe cell-specific localization of SMN protein.

**Figure 6:**
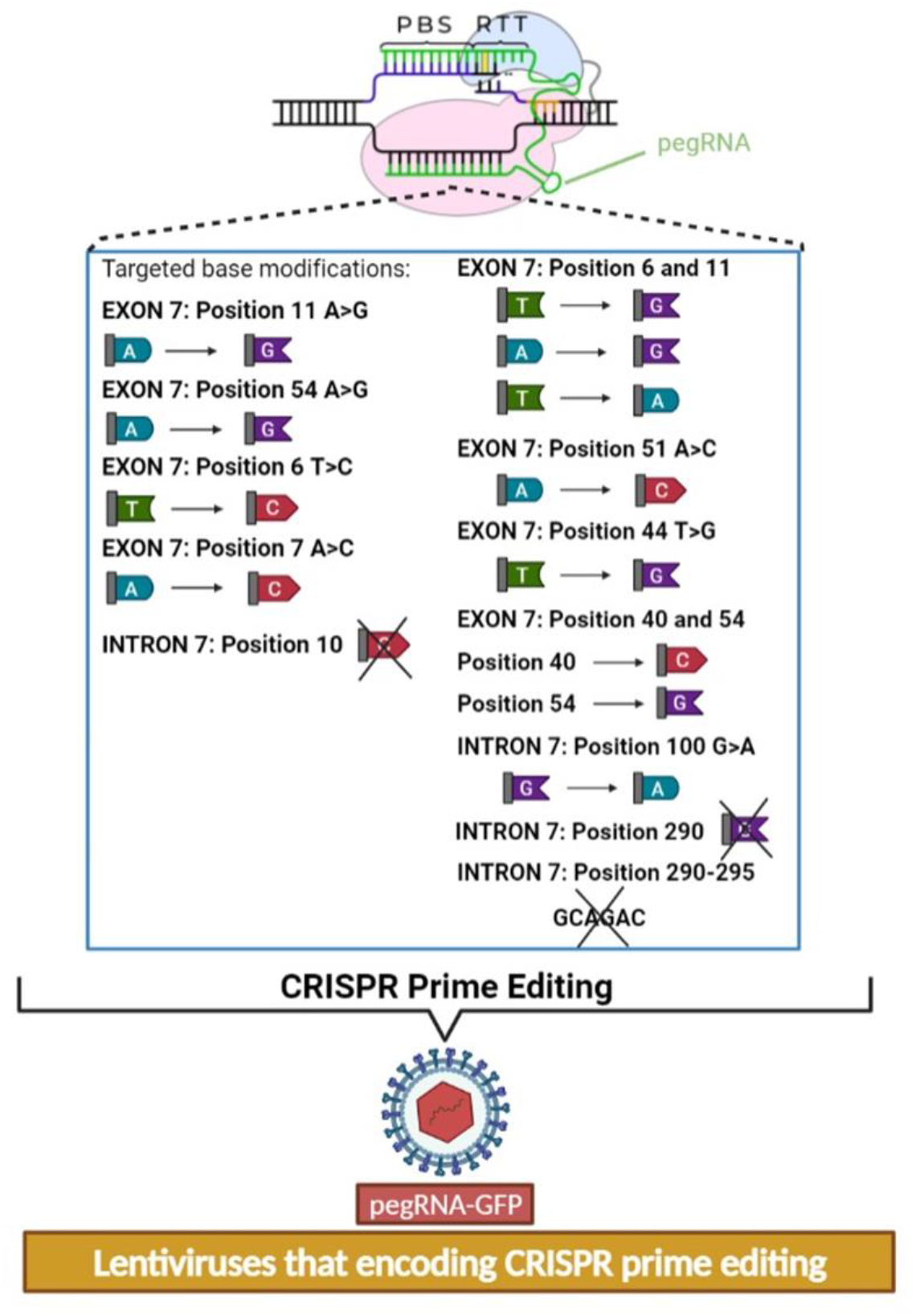
The CRISPR-PE screening assay with newly designed pegRNAs targeting new positions to perform the base editing.

## Discussion

Spinal Muscular Atrophy (SMA) is a fatal neuromuscular disease characterized by motor neuron loss and advanced muscle weakness, which occurs in functional SMN (Survival Motor Neuron) protein deficiency, which acts as a molecular chaperone in the formation of the spliceosome complex, which catalyzes the splicing of pre-mRNA, enabling mRNAs and non-coding RNAs to mature. Zolgensma, Nursinersen, and Evrysdi from SMA drugs; increase functional SMN protein expression using various gene therapy strategies, such as transgene delivery with viral vectors, Antisense oligonucleotide (ASO) technology, and chemical molecules that manipulate pre-mRNA processing, respectively. Although these drugs, which are included in the scope of genetic therapy, cannot be supplied by families due to their high prices, they do not offer a sustainable treatment opportunity for SMA as they cannot keep the functional SMN protein expression level stable in the long term (Story, 2011; Ramos and Chamberlain, 2015; Wang et al., 2020).

The CRISPR Prime Editing system has been frequently studied recently as a promising personal therapy method (Cohen, 2019; Y. Liu et al., 2020). It is predicted to correct genetic defects and correct approximately 89% of existing pathogenic variants. With this preliminary study, it has been shown for the first time in SMA disease that SMN protein expression can be increased by *SMN2* gene-targeted single nucleotide changes with CRISPR PE approaches. We evaluated the effect of our pegRNA systems on the SMN expression level in cells by transducing our pegRNAs into our CRISPR-PE encoding Jurkat cells. It was observed that the 840. C-T and 859. G-C changes targeted on the *SMN2* gene were successfully upregulated SMN expression in the cells. Especially, a combination of both modifications in pegRNA3 achieved the highest and most stable SMN expression for 14 days (**Figure 4**), showing that single nucleotide modification on the *SMN2* gene by CRISPR-PE approach can revert SMN expression in SMN-low cells. This showed that screening more pegRNA designs targeting other nucleotides on the *SMN2* gene (**Figure 6**) may provide SMN protein production at a physiological rate.

This study aimed to ensure stable and physiological SMN expression by using the CRISPR-PE system that will screen with 18 different pegRNA sequences in total. We will perform permanent base changes in a total of 18 regions in the *SMN2* gene, taking into account the *SMN2* gene exon 7/intron 7 specific variation analyses and mutagenesis studies (Cartegni and Krainer, 2002; Kashima and Manley, 2003; Singh et al., 2007; Hua et al., 2008; Gladman ve Chandler, 2009; Singh et al., 2013). Here, we investigated the effectiveness of the CRISPR-PE approach as an alternative to AAV-based Zolgensma, which transfers the functional *SMN1* gene, and ASO-based Spinraza, which provides transient SMN expression, for which we have applied for a patent (2021/018884), and functional and physiological SMN protein production and long-term. Next, we will plan to determine SMN protein level, stability, toxicity of the editing, tumorigenicity, and localization in motor neuron cells in a long time manner. Previous studies have shown that single point mutations on the *SMN2* gene can lead to an increase in SMN expression. However, until now, it has not been shown with the CRISPR system whether these mutations can be created synthetically in the laboratory and have the capacity to increase the production of SMN protein (Miccio et al., 2022). We designed a proof-of-concept experiment for the first time with this study by performing pegRNA designs that can create these point mutations, and we used the Jurkat cell line with low SMN expression for this. With this preliminary study, we showed that *SMN2* gene-targeted single nucleotide changes can increase SMN expression with the CRISPR-PE approach. Interestingly, although pegRNA2-encoding cells from our persistent CRISPR-PE and pegRNA-encoding Jurkat cell lines increased SMN expression on the 3rd day, it was found that the cells started to die when followed up to the 10th day. It is known that when the SMN gene is completely inactive, it is fatal in SMA type 0 patients and they die before they are born (Rao et al., 2018). Therefore, we speculate that expression of the pegRNA2 design in cells for 10 days may completely inactivate all copies of the SMN gene, and thus cells begin to die in a short time. In addition, we need to perform genetic sequencing to answer the questions of whether targeted genetic rearrangements have been achieved in PE-encoding cells in which all three pegRNA sequences are expressed, as well as whether off-target mutations have occurred. On the other hand, in these preliminary studies, we saw that SMN expression can turn completely negative as the cells begin to die. When we analyzed the same cells with both trypan blue and SMN monoclonal antibody, we found that the viable cell ratios were similar (data not shown). These results have shown us in many repeated experiments that SMN expression is also directly related to the viability of cells. In fact, the previous studies have reported that SMN protein might be related to Bcl-2 protein, which may result in affecting cell death pathway, namely apoptosis (Iwahashi et al., 1997; Coovert et al., 2000).

This study, in which we showed that genetic modifications that can cause an increase in SMN expression can be made with the CRISPR-PE system in Jurkat cells with low SMN expression, also allowed us to conduct extensive target mutation screening with the CRISPR-PE approach. Following this study, we saw that we could use pegRNA designs that we have proven to be effective in primary cells or neuronal cells isolated from patients with SMA. Therefore, in this study, we determined that we can increase SMN expression to physiological rates in primary fibroblast cells taken from SMA type1 patients. On the other hand, we measured SMN expression levels in many cell lines to test our CRISPR-PE system in neuron cells with our effective pegRNA designs, and we decided that we could use the A172 cell line in our future activity experiments. With this preliminary work, we have demonstrated that the CRISPR Prime Editing system is a very promising technology that could move into clinical applications in SMA disease. With this preliminary study, we have shown that the CRISPR Prime Editing method can be used more and more among patient-specific genetic therapy methods recently, where personal treatments have gained importance.

## Acknowledgment

We thank Ümit Fırat, Co-founder of A1 Life Sciences, Istanbul, Turkey for their sponsorship to accomplish the study. The project was selected as the best genetic therapy project in the competition of Rare Disease Challenge (RaDiChal) 2020. We thank Prof. Dr. Sevim Isik at Üsküdar University Stem Cell Production Application and Research Center (ÜSKÖKMER), Servet Tunoglu, MSc and Melisa Beyhan Yilmaz for the gift of A172, U87 and Hela cell lines. We thank Prof. Dr. Kasif Nevzat Tarhan, Founder-Rector of Üsküdar University, Istanbul, Turkey for his vision and support for genetic therapy field by founding TRGENMER Laboratory units in Üsküdar University.

